# Growth-supporting reductive defluorination by an amino acid-fermenting bacterial isolate

**DOI:** 10.64898/2026.11.15.766806

**Authors:** Weiyang Zhao, Qin Dong, Yiwen Zhu, Fengjun Xu, Yuan Fang, Bosen Jin, Rachel Lu, Shanmukh Bachhu, Patrick Moon, Ran Mei, Lawerence P. Wackett, Yujie Men

## Abstract

Per- and polyfluoroalkyl substances (PFAS), also known as “forever chemicals”, are ubiquitous environmental pollutants, highly persistent and toxic to humans and ecosystems. In addition to legacy PFAS, largely alkanoic acids, perfluorinated olefins and their corresponding enoic acids are emerging PFAS of concern. However, reported PFAS biodegradation is typically not growth-supporting, hindering the selection for effective defluorinating bacteria. Here, we report a new amino acid-fermenting bacterial isolate, *Sedimentibacter* sp. strain ZM-1, that performs high-rate reductive defluorination of a perfluoroenoic acid at physiologically relevant levels (mM). This process substantially enhanced energy conservation and cell growth, with three times higher growth than fermenting L-arginine alone and 50% higher than Stickland fermentation. The reductive defluorination redirected excess reducing equivalents (NADH) away from reducing protons and internal fermentation intermediates, as evidenced by decreased H_2_ production and downregulation of several NADH-dependent electron-bifurcating oxidoreductases. This defluorinating electron shunting drove arginine fermentation and oxidation of intermediates, such as ornithine and alanine, to form more acetate and CO_2_, leading to higher substrate-level ATP synthesis per arginine consumed. This process also generated reduced ferredoxin available for cell synthesis or anaerobic respiration, as genes involved in these processes were significantly upregulated. ZM-1 has a unique and effective CLC^F^-type fluoride exporter that works with the highly efficient defluorinating system to enable fluorotrophic growth. Since amino acid-fermenting anaerobes are common in the environment, such fluorotrophy opens new paths for natural selection of defluorination to drive natural and laboratory evolution of fluorophilic bacteria.

## Introduction

The unique physicochemical properties of C–F bonds contained in organofluorines make them a two-edged sword. On the one hand, their high resistance to moisture and harsh conditions has made per- and polyfluoroalkyl substances (PFAS) indispensable in many applications, including aerospace, semiconductors, data storage, and medical devices, since their invention in the 1940s^1,2^. Conversely, they become persistent (also known as “forever chemicals”) once released into the environment and can accumulate in organisms, causing adverse effects including carcinogenesis^1^. While most remediation currently focuses on legacy PFAS such as perfluorocarboxylic and perfluorosulfonic acids, perfluorinated olefins are emerging as the next big focus. They have traditionally been used in producing perfluorinated polymers such as Teflon and fluorinated ethylene propylene copolymers. More recently, perfluorinated olefin use has increased exponentially due to the expansion in data centers. Indeed, since 2019, > 60% of PFAS has gone toward data center applications^3–5^. There, they are used to cool high-density data center networks. Perfluorinated olefins are also seeing increasing use as solvents in electric vehicle battery packs and as blowing agents in producing polyurethane and polystyrene foams. In the pharmaceutical industry, they are used as fluorinated synthons to introduce fluorinated functional groups. Hydrofluoroolefins are widely used as heat-transfer agents in many industrial applications due to their efficiency and near-zero ozone-depletion potential^6^. Over time, they transfer into the atmosphere, where they may be converted into fluorinated carboxylic or enoic acid forms that could be carried down in precipitation^7^.

Microbial defluorination has rarely been observed in nature, especially for perfluorinated compounds, not only because of the high energy required to break C–F bonds, but also because microbes have not had sufficient time for evolutionary adaptation to occur^8,9^. Moreover, the highly oxidized carbon in per- and polyfluorinated compounds makes them poor electron donors^8^. They are, in theory, thermodynamically favorable as electron acceptors, similar to their chlorinated counterparts^10^. While reductive defluorination of fluorinated benzene rings is known, the process requires ATP and, as such, is not beneficial for the bacterium^11^. Similarly, other reports of microbial defluorination failed to show a connection to energetics for cell growth^12–14^. This is a critical roadblock, limiting selection for effective organofluorine degraders, as growth benefit is a key driver of evolution^15^. Here, we discovered a novel anaerobic bacterium that can utilize a perfluorinated enoic acid, one of the very few organofluorines that undergo microbial reductive defluorination, as an electron acceptor coupled to energy conservation. This exemplifies dehalorespiration of perfluorinated compounds and sets the first step toward discovering and designing more fluorophilic microbes that can benefit the environment and society.

### Fast reductive defluorination with high fluorinated compound tolerance by a novel environmental bacterial isolate, *Sedimentibacter* sp. strain ZM-1, possessing a unique CLC^F^-type fluoride exporter

An amino-acid-utilizing anaerobe was isolated from a reductive defluorinating anaerobic community originating from soil samples collected at a PFAS-impacted groundwater site and enriched by providing lactate and a perfluoroenoic acid, perfluoro(4-methylpent-2-enoic acid) (denoted “PFMeUPA”). After more than 15 rounds of subculturing, the enrichment culture achieved reductive defluorination of 100 µM PFMeUPA within three days (**Fig. S1**), much faster than any other reported cultures^14,16,17^. We performed anaerobic isolation by serial dilution and inoculation into semi-solid agarose tubes. Colonies exhibiting PFMeUPA-defluorinating activity in broth medium were subject to the next round of isolation until a highly enriched colony was obtained. It contained two metagenome-assembled genomes (MAGs) that were closest to *Clostridium homopropionicum* and *Sedimentibacter* sp, respectively. Since the same *Clostridium*-like MAG dominated (> 80%) in another highly enriched non-defluorinating colony, the *Sedimentibacter*-like MAG was then most likely responsible for PFMeUPA defluorination. Given that known *Sedimentibacter* species can grow on amino acids, whereas *Clostridium homopropionicum* cannot, we conducted further isolation using arginine and glycine as the primary energy and carbon sources, following a general *Sedimentibacter* recipe (see the Methods section). After several rounds of transfer in a liquid medium and isolation on semi-agar tubes, we obtained a *Sedimentibacter* isolate, which we named *Sedimentibacter* sp. strain ZM-1.

ZM-1 is a strictly anaerobic, Gram-positive, long-rod-shaped, amino-acid-utilizing bacterium (**Fig. 1A**). Similar to other *Sedimentibacter* isolates^18,19^, ZM-1 requires yeast extract as an essential growth supplement. Cell growth on amino acids became substantially lower when yeast extract was absent or added below 0.3% (w/v), even with all 20 amino acids provided. Thus, ZM-1 was maintained in basal medium supplied with arginine and glycine at 10 mM each, and 0.3% yeast extract. Considering the minor background growth on 0.3% yeast extract, ZM-1 showed additional fermentative growth with 10 mM individual amino acids, including arginine, glycine, serine, threonine, cysteine, methionine, histidine, and lysine. Other amino acids did not support cell growth above the yeast-extract-only background, although they were partially consumed when the yeast extract was provided alone (**Fig. S2**).

**Figure 1.**
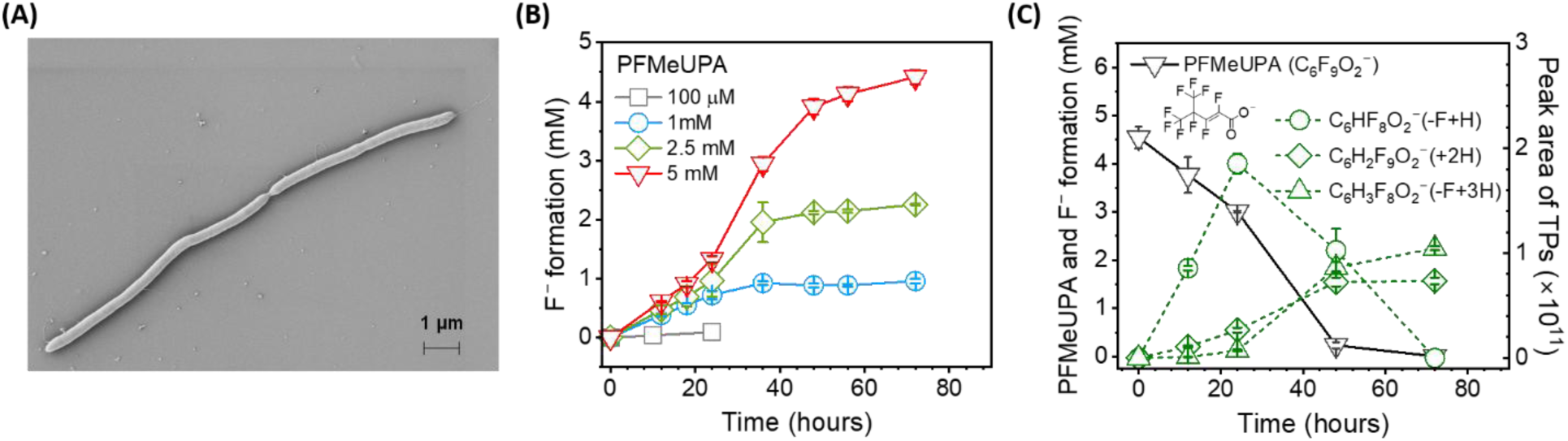
High-rate and high-tolerance reductive defluorination of PFMeUPA by *Sedimentibacter* sp. strain ZM-1. (A: scanning electron microscopy of ZM-1; B: fluoride formation from various initial PFMeUPA concentrations in ZM-1 supplied with 10 mM arginine and 10 mM glycine; C: the removal of 5 mM PFMeUPA and transformation product formation. Reductive defluorination product: C_6_HF_8_O_2_^−^, (CF_3_)_2_CFCF=CHCOO^−^. Hydrogenation product of PFMeUPA: C_6_H_2_F_9_O_2_^−^, (CF_3_)_2_CFCHFCHFCOO^−^. Hydrogenation product of C_6_HF_8_O_2_^−^: C_6_H_3_F_8_O_2_^−^, (CF_3_)_2_CFCFHCH_2_COO^−^).

Distinct from all other PFMeUPA-defluorinating cultures^14,16^, ZM-1 exhibited extremely fast reductive defluorination. At an initial PFMeUPA concentration of 100 µM, ZM-1 completed defluorination within 24 hours and within 72 hours for a much higher PFMeUPA concentration of 5 mM (**Fig. 1B**). Defluorination slowed down for subsequent doses of 5 mM PFMeUPA, with a maximum fluoride formation of 8 mM and incomplete removal of the third dose (**Fig. S3**). This could be due to increased cell toxicity of fluoride ions as they accumulated in the culture suspension. ZM-1 can also defluorinate PFMeUPA at lower concentrations (100 nM) to 4.7 nM within a 10-day incubation (**Fig. S4**). The transformation products of PFMeUPA in ZM-1 are the same as we previously reported for the PFMeUPA-defluorinating *Acetobacterium* species^14^, with reductive defluorination (the –F+H product in **Fig. 1C**) as the major pathway and hydrogenation (forming the other two transformation products **in Fig. 1C**) as the minor one.

Whole-genome sequencing by Illumina yielded a near-complete (94.87% completeness) ZM-1 genome of 4.05 Mbp (see detailed genome features in **Table S2**; GenBank accession No. JCAUJZ000000000). It has an average nucleotide identity (ANI) of 85.02% (63.8% alignment fraction) to the closest taxonomic placement, an unclassified *Sedimentibacter* species (GenBank accession No. GCA032065965) (**Fig. S5**). Thus, ZM-1 represents a new *Sedimentibacter* species. We also obtained the available pure cultures of three other *Sedimentibacter* species (i.e., *S. saalensis*, *S. acidaminivorans*, and *S. hydroxybenzoicus*) isolated from natural environments and examined their PFMeUPA defluorination activities. Interestingly, none of them exhibited defluorination kinetics comparable to ZM-1 (**Fig. S6A**). *S. saalensis* completed the defluorination of 100 µM after two weeks (**Fig. S6A**), much more slowly than ZM-1 (< 24 hours, **Fig. 1B**), while the other two strains only released a very low level of fluoride (**Fig. S6A**). This could be attributed to the unique functional genetics of ZM-1, including fluoride exporters that detoxify released fluoride ions and are essential for defluorination to occur inside cells^14,20^.

Bacteria typically use two types of fluoride exporters to mitigate cell toxicity: the CLC^F^-type F^−^/H^+^ antiporter and the CrcB-type selective F^−^ channel^21^. Recent studies showed that the CLC^F^-type F^−^/H^+^ antiporter is advantageous over the CrcB-type exporter in coping with extremely high fluoride stress (above mM levels)^22^. ZM-1 genome possesses two annotated CLC^F^-type F^−^/H^+^ antiporters. The first one, CLC^F^-1, shares a highly similar amino acid sequence identity (79%) with that in *S. saalensis*, which showed complete transformation of PFMeUPA but at a much slower rate (**Fig. S6A**). In contrast, the second one, CLC^F^-2, has only 34 – 40% amino acid identity to CLC^F^-1 and other previously confirmed CLC^F^-type F^−^/H^+^ antiporters^23^ (**Fig. S6B**). Moreover, the gene (*eriC*) encoding CLC^F^-2, rather than CLC^F^-1, was significantly upregulated (> 64-fold change) in actively defluorinating ZM-1 and in ZM-1 exposed to 3 mM NaF (**Fig. S7** and **Table S3**), indicating that CLC^F^-2 is the functional fluoride exporter under high fluoride stress. This *eriC* gene was also among the most up-regulated genes under the PFMeUPA-defluorinating condition (**Table S3**). The rapid defluorination and fluoride secretion via the CLC^F^ F^−^/H^+^ antiporter may cause instant local acidification, while hydrogenation reactions and amino acid metabolism may consume H^+^ and form ammonia to neutralize H^+^, respectively. Overall, there was only a slight decrease of pH in ZM-1 with PFMeUPA (**Table S4**). Genes encoding several Na^+^/H^+^ antiporters and a K^+^-uptake transporter (KtrAB), which were important for maintaining H^+^ homeostasis^24^, were upregulated active defluorination (**Table S3**).

Meanwhile, the cell-free protein extract of ZM-1 pre-grown in PFMeUPA-free medium revealed much faster defluorination of PFMeUPA in *in vitro* assays than the lysate of other *Sedimentibacter* species (**Fig. S8**). This suggested that the higher rate of reductive defluorination by ZM-1 cells was not only due to the effective fluoride detoxification enabled by the unique CLC^F^-type F^−^/H^+^ antiporter but rather to the intrinsic rate of the defluorination reaction. Protein extracts were prepared using a detergent lysis procedure that can support the activity of soluble or membrane proteins, leaving the localization of defluorinase activity open to further investigation.

### Enhanced cell growth with the perfluoro-enoic acid as an electron acceptor for redox balance and energy conservation

The physiologically relevant (mM) levels of fluorinated compounds that ZM-1 can handle led to the hypothesis that reductive defluorination could play a direct role in ZM-1 cellular metabolism, likely serving as an electron acceptor for energy conservation. To test this hypothesis and examine the potential growth benefit of carrying out high-level reductive defluorination, we compared ZM-1 growth under the two conditions: (i) fermentation on individual amino acids and (ii) with PFMeUPA as an external electron acceptor. Cells were pre-grown on arginine and glycine and washed three times in anaerobic mineral salts medium before inoculation to avoid carryover of organic substrates. Notably, the growth of ZM-1 was substantially stimulated in the presence of PFMeUPA coupled with individual amino acids (arginine, alanine, ornithine, glycine, and threonine), as well as 0.3% yeast extract containing a mixture of amino acids, compared with growth without PFMeUPA (**Fig. 2**). Yeast extract alone gave marginal growth, but with PFMeUPA, the growth was stimulated by one order of magnitude. When individual amino acids were supplied, growth with arginine plus PFMeUPA was more than three times higher than with arginine alone and 50% greater than with arginine and glycine, which is considered a superior fermentation condition for amino acid-utilizing anaerobes via Stickland reactions. PFMeUPA also stimulated growth, but to a lesser degree, for other fermentable amino acids. Interestingly, alanine or ornithine alone did not support growth, but with PFMeUPA, growth was significant, and alanine or ornithine utilization increased (**Fig. S9**).

**Figure 2.**
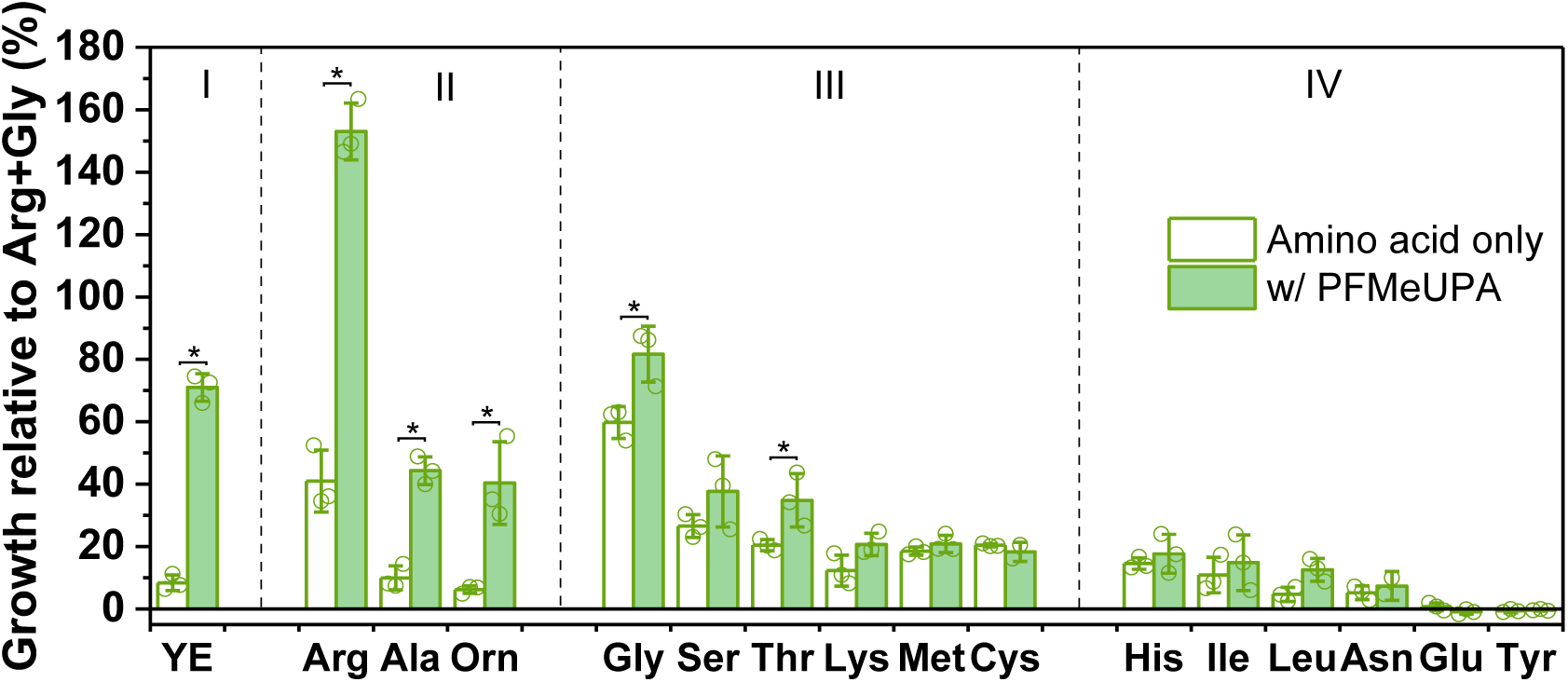
Relative cell growth of ZM-1 on individual amino acids with and without PFMeUPA to that on arginine and glycine (amino acids were provided at 10 mM each; I: yeast extract (YE) control, 0.3% w/v; II: arginine and related degradation intermediates, alanine and ornithine (Orn); III: other fermentable amino acids; IV: amino acids showed consumption in YE-only control but did not support cell growth when provided individually in excess; cells were quantified by total genomic DNA at the highest OD_600_; *: significant difference in cell growth with PFMeUPA compared to without, Student’s t-test, *p* < 0.05, n = 3).

We then focused on the arginine and PFMeUPA condition, which exhibited the most growth benefit, with the arginine-only condition as the control. Cells grown on arginine and PFMeUPA started to outgrow those without PFMeUPA, as defluorination reached its highest rate after one day (**Fig. 3A**). While arginine consumption was similar (∼ 8 mM after 4 days), the metabolite profiles differed significantly. With PFMeUPA, acetate and CO_2_ accumulated to substantially higher levels (∼13 mM and ∼ 35 mM, respectively) (**Fig. 3B & C**). Consistently, more citrulline, ornithine, and alanine were consumed with PFMeUPA than without (**Fig. S10**). This indicates deeper oxidation of arginine and fermentation intermediates, driven by reductive defluorination of PFMeUPA, which consumes electrons and helps the redox balance of the cell. H_2_ was produced during arginine fermentation to balance cellular redox, and H_2_ production decreased significantly when cells were grown with PFMeUPA (**Fig. 3C**), indicating electron shunting toward PFMeUPA reduction. We noted that acetate and CO_2_ levels were higher than could be accounted for by the additional consumption of citrulline, ornithine, and alanine (**Fig. S10**). This could be attributed to enhanced fermentation and oxidation of amino acids, such as arginine and alanine, contained in the 0.3% yeast extract required in the medium (**Fig. S1**).

**Figure 3.**
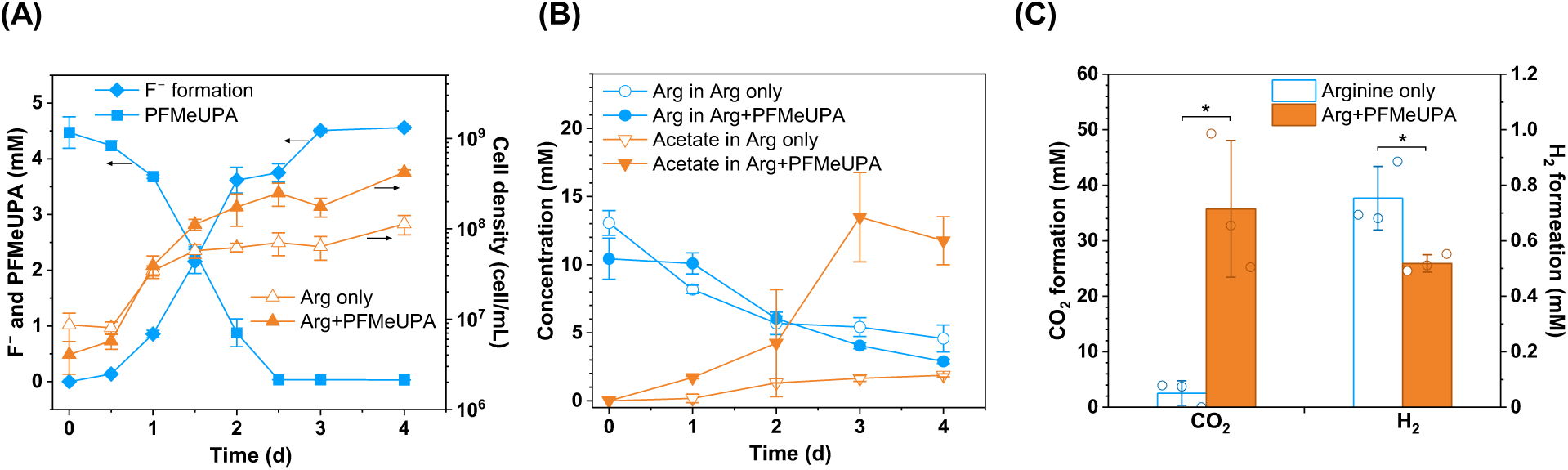
Comparison of ZM-1 metabolic activities when grown on arginine (10 mM) without and with PFMeUPA (5 mM) (A: F^−^ formation, PFMeUPA decay, and cell growth. RNA samples were taken on Day 1.5; B: arginine utilization and acetate formation; C: CO_2_ and H_2_ measured in a repeated batch under the same growth conditions; CO_2_ and H_2_ formation was calculated based on gaseous measurement, headspace pressure, and pH shown in **Table S4**; *significant difference between with and without PFMeUPA conditions by Student’s t-test, *p* < 0.05, n=3; see **Fig. S10** for end-point profiles of other metabolites and cell growth in this repeated batch).

ZM-1 possesses two arginine utilization pathways. The arginine deiminase (ADI) pathway (**Reactions 1 – 3** in **Fig. 4**) generates one ATP via substrate-level phosphorylation via citrulline, with ornithine and CO_2_ as the products. This process is redox neutral without formation of reducing intermediates like NADH. Another is the arginase pathway (**Reaction 4, Fig. 4)**, forming ornithine. ZM-1 can further oxidize ornithine to alanine and acetate, and alanine to acetate and CO_2_ (**Reactions 5 – 9** and **Reactions 10 – 12**, respectively, in **Fig. 4**). This leads to the formation of another ATP along with reducing molecules including NADH and reduced ferredoxin (Fd_red_). Without external electron acceptors, cells must divert the excess reducing power to internal electron acceptors, such as protons and intermediates from arginine fermentation, to maintain cellular redox balance. ZM-1 genome lacks proline reductases and does not appear to reduce proline to 5-aminovalerate, as reported in *Clostridium sticklandii* (reclassified as *Acetoanarobium sticklandii*)^25^, to maintain redox balance. Nonetheless, ZM-1 expressed several hydrogenases, including electron-bifurcating hydrogenase complexes (HydABCD), which can produce H_2_ to re-oxidize NADH (**Fig. 4, Table S3**). Although we did not detect common reduced fermentation products, such as propionate and butyrate, we did detect high expression of genes encoding NADH-dependent electron-bifurcating acyl-CoA dehydrogenase complexes and fumarate/urocanate reductase-like flavocytochrome c that may consume NADH to form reduced products (**Fig. 4**, **Table S6**). These reduction processes made less use of the electrons from arginine, resulting in less ATP and cell growth under the arginine-only condition. In contrast, 5 mM PFMeUPA, as an alternative electron acceptor, redistributed electron flow, leading to more ATP formation per arginine consumed, thereby higher cell growth. PFMeUPA may play a role similar to 10 mM glycine in Stickland fermentation, but it is notably a better electron acceptor for energetics, supporting 50% higher cell growth than Stickland fermentation (**Fig. 2**). One may also not exclude the possibility of the coupling of reductive defluorination to anaerobic respiration (i.e., oxidative phosphorylation) via the Rnf complex in ZM-1 (see the next section). More significantly, regardless of how additional ATP is synthesized, this sets up a new fluorotrophic metabolism involving the breakdown of perfluorinated compounds.

**Figure 4.**
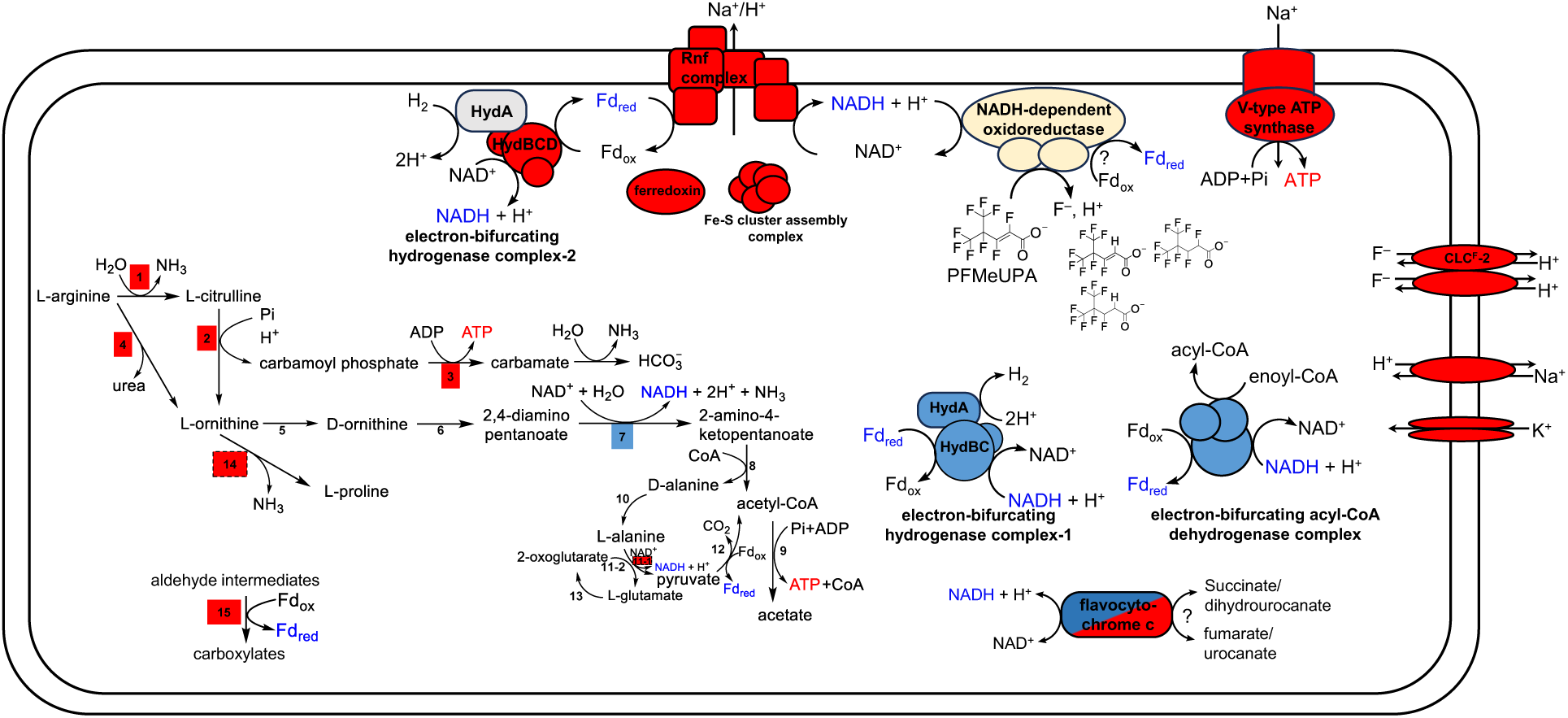
Proposed catabolic degradation of arginine coupled to dehalorespiration of PFMeUPA for energy conservation (red shade: up-regulation, blue shade: down-regulation, adj. *p* < 0.01, > 1.5-fold change, and grey shade: no significant transcriptional change; all genes were expressed with a normalized transcript count > 100, and genes with normalized transcript counts < 500 are marked in dashed boxes, samples taken on day 1.5 in **Fig. 3**; yellow shade: potential defluorinases; ATP synthesis in red text; reducing power production in blue text; enzymes involved in the arginine catabolism pathway: 1. arginine deiminase; 2. ornithine carbamoyltransferase; 3. carbamate kinase; 4. arginase; 5. ornithine/alanine racemase; 6. D-ornithine 4,5-aminomutase; 7. 2,4-diaminopentanoate dehydrogenase; 8. 2-amino-4-ketopentanoate thiolase; 9. acetate kinase; 10. alanine racemase; 11-1. alanine dehydrogenase; 11-2. alanine aminotransferase; 12. pyruvate: ferredoxin oxidoreductase; 13. glutamate dehydrogenase; 14. ornithine cyclodeaminase; 15. tungsten-containing aldehyde: ferredoxin oxidoreductase.

### Transcriptional indicators of enhanced arginine oxidation and anaerobic respiration with PFMeUPA

In this analysis, we focused on regulated genes specifically in response to the fluorophilic metabolism by excluding those differentially expressed in ZM-1 when exposed to NaF (**Table S3**). Consistent with physiological observations, transcriptomes revealed significant upregulation of genes involved in arginine catabolism, oxidative energy conservation, and gluconeogenesis when grown on arginine and PFMeUPA compared with arginine alone (**Fig. 4**, **Table S3, Table S5– 7**). Genes involved in both ADI and Arginase pathways, as well as downstream oxidation pathways, were significantly upregulated (adj. *p* < 0.01, **Table S5**), in line with the increased consumption of citrulline, ornithine, and alanine. Additionally, some of the aforementioned redox-balancing electron-bifurcating hydrogenase, butanoyl-CoA reductase complexes, and flavocytochrome c (fumarate reductases) were significantly downregulated (**Fig. 4** and **Table S3**). This corroborates the decreased H_2_ production and again suggests redistribution of electron flow (as NADH) toward PFMeUPA. ZM-1 lacks electron transport complexes or previously identified reductive dehalogenases for reductive dechlorination. It is thus reasonable to infer that PFMeUPA was reduced directly by an NADH-dependent oxidoreductase (**Fig. 4**).

Furthermore, genes encoding the Rnf complex and the V-type ATP synthase were significantly upregulated (adjusted *p* < 0.01), as were genes encoding ferredoxin and Fe-S cluster assembly proteins, which are relevant components of the Rnf complex (**Fig. 4** and **Table S6**). The Rnf complex plays an essential role in anaerobic respiration in other anaerobic bacterial species^26^, including the homoacetogen *Acetobacterium woodii*^27^ and the amino-acid-utilizing *Acetoanarobium sticklandii*^25^ (former name: *Clostridium sticklandii*), pumping Na^+^/H^+^ out of the cytoplasmic membrane driven by the oxidation of Fd_red_ ^25,26,28^. These anaerobic bacteria also possess V-type ATP synthase, which can carry out oxidative phosphorylation driven by the sodium motive force (*smf*) created by the Rnf complex. Reversibly, the ATP synthase can translocate Na^+^ outside of the membrane driven by ATP hydrolysis, then the *smf* drives Fd_ox_ reduction with NADH at the Rnf complex^26^. This could be used to regenerate Fd_red_ needed for various metabolic processes and re-oxidize NADH to NAD^+^. The upregulation of Rnf complex and V-type ATP synthase operons in ZM-1 with PFMeUPA may be caused by increased activity in either direction. On the one hand, deeper oxidation of ornithine and alanine produces more Fd_red_ (**Fig. 4**), which could drive the forward Rnf reaction to conserve energy via oxidative phosphorylation. This complements the enhanced substrate-level ATP formation from fermentation to benefit cell growth. On the other hand, elevated ATP could be used in the reverse reaction of the Rnf complex, forming Fd_red_, which may be needed for anabolism. For example, the highly expressed pyruvate:ferredoxin oxidoreductase (**Table S3** & **S5**) could perform the reverse reaction using Fd_red_ to produce pyruvate, a precursor of gluconeogenesis. The gluconeogenic pathway is essential for cell synthesis in anaerobic bacteria grown on non-sugar substrates, such as amino acids. During the exponential growth phase of ZM-1 with PFMeUPA, the entire gluconeogenesis pathway was significantly upregulated by more than two-fold (**Table S7**). Notably, one electron-bifurcating hydrogenase complex (HydBCD/NuoFEG) and one Tungsten-containing aldehyde:ferredoxin oxidoreductase (**Fig. 4**, **Table S5 & S6**) were upregulated. Given the decreased H_2_ level and electron shunting, it is reasonable to infer that these upregulated genes were involved in the oxidation direction, consuming H_2_ and aldehyde intermediates and forming Fd_red_ to supplement the Fd_red_ pool, as it is being utilized for both energy conservation and biosynthesis. Additionally, the defluorinating enzyme in ZM-1 might also couple to Fd_red_ formation in an electron-bifurcating complex similar to what was implied in *Acetobacterium* spp.^14^. What is different from the electron-bifurcating caffeate reduction complex in *Acetobacterium* spp. could be the much higher enzyme affinity and kinetics, and the no requirement for CoA activation, which would make the reductive defluorination in ZM-1 more energy-efficient.

### Implications in the discovery and design of microorganisms defluorinating highly fluorinated compounds

It has long been asked whether microorganisms can derive a selective benefit from transforming highly fluorinated compounds such as PFAS. Some researchers suggest that fluorine is intrinsically harmful to any possible life forms^29^, while others suggest that planets with fluorine-dependent life are plausible^30^. Most of Earth’s biosphere does not use fluorine in its metabolism^31^. One rare exception is the production of monofluorinated toxins such as monofluoroacetate (MFA), which certain plants release to kill bacterial pathogens. Over a long time, bacteria have evolved mechanisms to cope with MFA. The most common one is via hydrolytic defluorination catalyzed by 2-haloacid dehalogenases, which forms the non-toxic glycolate that enters central metabolism. Here, although MFA is not directly involved in energetics, the detoxification of MFA and the additional growth benefit provided by the product were the driving force of this evolution. Similarly, the degradation of other mono- or less fluorinated compounds (e.g., with one trifluoromethyl group, –CF_3_) is coupled with the formation of non-fluorinated products that may serve as energy and carbon sources for cell growth^32,33^. This is much less likely with highly fluorinated compounds (e.g., PFAS), where most carbons are fluorinated. A more likely way for perfluorinated compounds to be involved in metabolism is by serving as electron acceptors in energetics. However, the literature lacks evidence for positive selective pressure operating on the energy metabolism of per- and polyfluorinated compounds, although growth-supporting defluorination of MFA via reductive processes was reported in one anaerobic bacterium (*Cloacibacillus porcorum* strain MFA1)^34^. Thus, creating metabolism to consume highly fluorinated compounds will require a ground-up redesign.

Here, we found promise for metabolic selection of microorganisms that break down highly fluorinated contaminants by discovering this new *Sedimentibacter* isolate ZM-1, which can conserve energy for growth through reductive defluorination of a perfluorinated compound. Although metagenomic analyses have detected *Sedimentibacter* species in communities carrying out reductive dechlorination of chlorinated solvents^35^, they remain understudied as isolates, especially their metabolic traits and the underlying molecular basis of energetics. This study fills this knowledge gap and underscores the critical yet previously overlooked role *Sedimentibacter* species may play environmentally in the metabolic transformation of organofluorines. Because ZM-1 was isolated from PFAS-impacted soils, this suggests adaptive evolution of microorganisms during long-term exposure to various PFAS compounds in the environment. Isolating, identifying, and characterizing “fluorophilic” microorganisms and enzymes can provide a blueprint for designing fluorine-dependent metabolism for biotechnological purposes experimentally and computationally, hence accelerating natural selection that would otherwise take much longer. Such fundamental research is pivotal and should continue not only because of the beauty of discovering Mother Nature’s diversity and new capabilities but also because biotechnologies using effective microbial degraders would still be a superior approach for *in situ* remediation. This is the first, yet essential, step toward an ultimate goal of transforming science into applicable biotechnologies. Although long and slow, this is the right path to take us there. It took decades from the first discovery of a dehalorespiring bacterium to a thorough understanding of the responsible enzyme for reductive dechlorination, eventually leading to real-world applications for chlorinated solvent remediation^36–39^. It may take much longer to fully identify and characterize the enzymes required for reductive defluorination of highly fluorinated compounds, given the completely different physicochemical properties. Roadblocks we foresee along the path include complex procedures for purification of oxygen-sensitive enzymes and a lack of established genetic modification tools for non-canonical anaerobic bacteria to optimize substrate specificity and defluorination extent. Nonetheless, we should remain faithful to overcoming these hurdles step by step, as our findings continue to shed light on the path, and we have not encountered a dead end.

## Materials and Methods

### Chemicals

(E)-perfluoro(4-methylpent-2-enoic acid) (PFMeUPA, CAS: 103229-89-6) was purchased from SynQuest Laboratories (Alachua, FL, USA) and used without further purification (see details in **Table S1**). Other general chemicals, including amino acids, were purchased in ACS grade (≥ 95% purity) from Sigma-Aldrich (St. Louis, MO, USA).

### Bacterial cultures and medium recipes

Besides ZM-1, which was isolated in this study, three other amino-acid-utilizing *Sedimentibacter* species used in this study, i.e., *S. saalensis* (DSM 13558), *S. acidaminivorans* (DSM 24004), and *S. hydroxybenzoicus* (DSM 7310), were obtained from German Collection of Microorganisms and Cell Cultures GmbH (DSMZ). *S. saalensis*, *S. acidaminivorans*, and *S. hydroxybenzoicus* were revived and maintained in their respective DSMZ Medium 943, 1616, and 643 (https://www.dsmz.de/), respectively, to achieve optimal growth before being subject to defluorination assays.

### Enrichment setup

Soil and groundwater samples were collected from different locations at a field site impacted by PFAS-containing aqueous film-forming foam. Two grams of soil from each location were weighed and inoculated into a 125-mL serum bottle containing 25 mL anaerobic basal medium, the same as previously used, and 25 mL filtered (0.22 µm) groundwater with the addition of 35 mM lactate and 100 µM PFMeUPA (filter-sterilized; the stock solutions were made in Milli-Q water), similar to conditions used before for defluorinating enrichments^16^. Microcosms were incubated at 30 °C in the dark without shaking. One soil microcosm exhibited fast defluorination activity after two weeks and was subsequently transferred (10% v/v) into the same medium more than four times before proceeding to anaerobic isolation.

### Anaerobic Isolation

The most active defluorinating enrichment culture was used to obtain defluorinating isolates. The anaerobic isolation was conducted in 25-mL sealed culture tubes containing 10 mL of the same medium used for the enrichment culture, supplemented with 0.7% (w/v) agarose to make semi-solid medium. After autoclaving, before the medium began to solidify, serial-diluted enrichment culture (10^−2^ – 10^−14^) was inoculated and supplied with 35 mM lactate and 100 µM PFMeUPA. The isolation tubes were inoculated at 30 °C in the dark without shaking. Colonies were picked up using a long needle and transferred into 10 mL of liquid medium containing lactate and PFMeUPA. Fluoride was measured over time. Tubes with the highest defluorination activity were subjected to the next round of isolation on semi-solid agarose medium. After several rounds of isolation, metagenomic sequencing was conducted for one highly defluorinating colony and another without any defluorination activity. The defluorinating colony consisted of two major bacterial species, *Clostridium homoproprionicum* and *Sedimentibacter* spp., and the former was also dominant in the non-defluorinating colony. Thus, further isolation from the highly enriched colony was performed using *Sedimentibacter*-specific medium (DSMZ Medium 643). After several rounds of isolation, we obtained a pure *Sedimentibacter* culture (ZM-1), whose purity was examined by microscopy and (meta)genomic sequencing, resulting in a single assembled *Sedimentibacter* genome (**Table S2**). ZM-1 was maintained in DSMZ Medium 643 containing arginine (10 mM), glycine (10 mM), and 0.3% (w/v) yeast extract.

### Whole-genome sequencing and analysis

ZM-1 cells were harvested by centrifugation at 8,000 × g for 15 min (4 °C), then stored at –80 °C for further use. Genomic DNA was extracted from cell pellets using a DNeasy Blood & Tissue Kit (Qiagen, Germantown, MD) according to the manufacturer’s instructions. The DNA was sent to SeqCenter (Pittsburgh, PA; https://www.seqcenter.com/service/illumina-dna-sequencing/) for whole-genome sequencing by Illumina NovaSeq (2 × 150 bp). *De novo* assembly was performed by SPAdes, and genes in the resulting contigs were annotated by RAST.

### Cell growth measurement by Qubit

Genomic DNA extracted from 1.5 mL cell culture was quantified by a Qubit 4 Fluorometer (Thermo Fisher Scientific) according to the manufacturer’s instructions. Cell densities (genome copies per mL) were then calculated from the total DNA concentration and the genome size of ZM-1 (∼4.05 Mbp) as described previously^40^.

### Reverse-transcription quantitative PCR (RT-qPCR)

ZM-1 cells were collected by centrifugation at 8,000 × g, 4 ℃, for 20 min. Total RNA was extracted using the TRIzol Max Bacterial RNA Isolation kit (Life Technologies, Carlsbad, CA) and cleaned with a Zymo RNA Clean and Concentrator kit (Zymo Research, Irvine, CA). The quality and concentration of RNA samples were measured on a NanoDrop spectrophotometer (Thermo Fisher Scientific, Waltham, MA) and a QuantStudio 3 Real-Time PCR System to ensure the absence of DNA. Reverse transcription of the purified RNA samples was then carried out using SuperScript III First-Strand Synthesis SuperMix (Thermo Fisher Scientific). The resulting cDNA was subjected to qPCR targeting the *eriC* gene (GeneID 1445 in ZM-1 genome, **Table S3**), which encodes a unique CLC^F^-type F^−^/H^+^ antiporter, with the housekeeping DNA gyrase gene *gyrB* as the control. The qPCR primers (5’→3’) for *eriC* were CTATTTGCCATAGGAGCCTC (forward) and ATCCGAAAACTGCCAGAAGC (reverse), and for *gyrB* were ATGATGTTGCTATAACCGGAG (forward) and TTGGTCTTTGTCTGCCCTTC (reverse). Each 20-μL reaction contained 2.5 μL cDNA, 10 μL 2× PowerUp SYBR Green Master Mix (Thermo Fisher Scientific), and 1.25 μL each of 10 μM forward and reverse primers. Thermal cycling conditions consisted of initial enzyme activation at 95°C for 2 min, followed by 40 cycles of denaturation at 95°C for 1 s and annealing/extension at 60°C for 30 s. ΔΔC_T_ was used to determine the fold change in *eriC* transcripts.

### RNA-sequencing and comparative transcriptomics

ZM-1 cells grown on arginine only and on arginine and PFMeUPA were collected after 1.5 days by centrifugation at 8,000 × g, 4 ℃, for 20 min. Total RNA was extracted using the method described in the above section. Quantified and quality-controlled RNA samples were sent to SeqCenter for RNA-sequencing by Illumina NovaSeq (2 × 150 bp). Demultiplexing, quality control, and adapter trimming were performed with bcl-convert (v4.2.4). Raw sequence data (.fastq files) were processed online with the Galaxy server (https://usegalaxy.org/). Sequence read quality was checked by FastQC^41^. Paired-end reads were joined with Fastq interlacer and mapped to ZM-1 genome (GenBank accession number: JCAUJZ000000000) using Bowtie2^42^. Bowtie2-generated BAM files were used to generate transcript count tables with FeatureCount^43^. Differential gene expression analysis between two conditions was performed with DESeq2^44^. Differentially expressed genes at the 99% confidence level (adj. *p* < 0.01) and normalized counts > 100 in at least two replicates under one condition were considered for further analysis.

### Analytical methods

The F^−^ concentration in the culture suspension was measured using an ion-selective electrode (ISE; HACH, Loveland, CO) connected to an HQ30D Portable Multi Meter (HACH). Fluorinated compounds and PFMeUPA transformation products were detected and quantified (if reference compounds are available) on an ultra-high-performance liquid chromatography coupled with high-resolution tandem mass spectrometry (UHPLC-HRMS/MS) (Orbitrap Exploris 240, Thermo Fisher Scientific) using the same method developed and reported previously^40,45^. For low PFMeUPA concentrations (i.e., an initial concentration of 100 nM), an LC triple-quad MS/MS (Agilent Technologies, La Jolla, CA) was used in MRM mode with a signature transition from m/z 275.0 to 231.0 at a fragmentation voltage (Frag) of 118.0 eV and a collision energy (CID) of 4 eV. The retention times and limits of quantification for the investigated compounds are listed in **Table S1**.

All amino acids were analyzed on the same LC-HRMS/MS (Orbitrap Exploris 240). A 2 μL sample was loaded onto an ACE AQ column (particle size 3 µm, 3 mm × 150 mm, Avantor, Radnor Township, PA) and eluted using Milli-Q water with 0.1% (v/v) formic acid (A) and HPLC-grade acetonitrile with 0.1% (v/v) formic acid (B) at a flow rate of 200 µL/min. A linear elution gradient was used: 100% A for 0 – 1.5 min, 100– 0% A for 1.5 – 12 min, 0% A for 12–13.5 min, and 100% A for 13.5 – 15 min. For the HRMS part, positive electrospray ionization was used for a full MS scan (m/z = 50 – 250) at a resolution of 240,000 @ m/z 200. Samples were appropriately diluted to fall within the calibration standard series range, 0.1 – 100 µM.

Other small fatty acids were measured by HPLC coupled with a photodiode array (PDA) detector (Shimadzu). A 10 µL sample was loaded onto a Bio-Rad Aminex HPX-87H ion-exclusion column heated to 60°C, eluted using 5 mM H_2_SO_4_ at a flow rate of 0.6 mL/min, and detected at 210 nm.

Headspace samples (1 mL) were analyzed using a GC-2030 gas chromatograph (Shimadzu, Kyoto, Japan) equipped with an SH-Q-BOND PLOT column, an SH-Msieve 5A PLOT column, a thermal conductivity detector (TCD), and a flame ionization detector (FID). Samples first passed through the SH-Q-BOND PLOT column, where CO₂ was separated from H₂, O₂, N₂, and CH₄. Before CO₂ elution, H₂, O₂, N₂, and CH₄ were directed to the SH-Msieve 5A PLOT column and detected by TCD; the effluent was subsequently routed to the FID for more sensitive CH₄ detection. CO₂ eluting later from the SH-Q-BOND PLOT column was directed to the FID channel through a time-programmed valve event. Samples were introduced through a split inlet at 150 °C using helium as the carrier gas. The oven was held at 35 °C for 4 min and then increased to 105 °C at 30 °C/min, giving a total run time of 7 min. The FID was operated at 400 °C with argon make-up gas, H₂, and air at 20, 32, and 200 mL/min, respectively. The TCD was operated at 150 °C and 75 mA, with argon as the make-up and reference gas at 15 and 45 mL/min, respectively. H₂ was quantified using external calibration standards.

## Data availability

ZM-1 genome was deposited in GenBank under the accession number JCAUJZ000000000. Transcriptomic data were deposited in the Sequence Read Archive (SRA) under the accession numbers SAMN61879765 – SAMN61879769. Preview link for reviewers: https://dataview.ncbi.nlm.nih.gov/object/PRJNA1498711?reviewer=o49gssr8ec6hb8uc5njlckgn5n. The sequences will be made publicly available upon the publication of the manuscript.

## Acknowledgements

This study was partly supported by the National Institute of Environmental Health Sciences (Award No. R01ES032668, 2021 – 2026) for W.Z. and Y.M., the Strategic Environmental Research and Development Program (ER20-1541 2020 – 2026) for Q.D., Y.Z., B.J., Y.M., and L.P.W., the John Babbage Endowed Chair Fund in the Bourns College of Engineering at the University of California, Riverside for F.X., and the internal fund for Q.D. and Y.M.

## Author Contributions

Conceptualization: Y.M. Experimental data collection: W.Z., Y.Z., Q.D., B.J., S.B., P.M., Y.F., and R.M. Data analysis: W.Z., Q.D., Y.Z., F.X., L.P.W., and Y.M. Writing: Y.M., W.Z., and Q.D. Review & Editing: W.Z., Q.D., Y.Z., F.X., Y.F., B.J., S.B., P.M., R.M., L.P.W., and Y.M. Supervision: Y.M.; Funding acquisition: L.P.W. and Y.M.

## Notes

### Competing Interest Statement

The authors have declared no competing interest.

