## Supplementary material for "Growth-supporting reductive defluorination by an amino acid-fermenting bacterial isolate": SI Table S1-2, 4-7, Figure S1-10

**This PDF file includes:**

Table S1 – S2, Table S4 – S7

Figure S1 – S10

**Other Supplementary Materials for this manuscript include the following:**

Supplementary Table: Excel file for Table S3

**Table S1** Chemical and analytical information of PFMeUPA and its transformation products

| Structure (Acronym) | CAS | Name (Formula) | Purity | $[M-H]^-$ | RT (min) | LOQ on LC-HRMS (μM) |
|---|---|---|---|---|---|---|
| 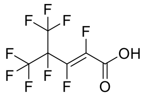 (PFMeUPA) | 103229-89-6 | 4-(Trifluoromethyl)hexafluoropent-2-enoic acid ($C_6HF_9O_2$) | 97% | 274.9754 | 6.37 | 0.5 (0.1 nM on LC-triple-quad MS) |
| 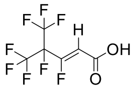 | - | Transformation product of PFMeUPA ($C_6H_2F_8O_2$) | - | 256.9854 | 7.80 | - |
| 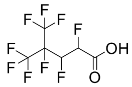 | - | Transformation product of PFMeUPA ($C_6H_3F_9O_2$) | - | 276.9917 | 6.83 | - |
| 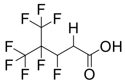 | - | Transformation product of PFMeUPA ($C_6H_4F_8O_2$) | - | 259.0011 | 6.92 | - |

**Table S2 Genomic features of *Sedimentibacter* sp. ZM-1**

| Features | |
|---|---|
| Genome completeness (%) | 94.87 |
| Contamination (%) | 2.45 |
| Genome size (Mbp) | 4.05 |
| G+C content (%) | 36.95 |
| Number of CDS | 3914 |
| Number of tRNAs | 55 |

**Table S4 $H_2$ and $CO_2$ measurement**

| Sample | Gauge pressure (pressure increase in headspace compared to the atmosphere) (psi) | $H_2$ (%) | $CO_2$ (%) | pH |
|---|---|---|---|---|
| **Arg-1** | 0.0 | 2.74% | 1.81% | 7.89 |
| **Arg-2** | 0.0 | 2.69% | 1.68% | 7.92 |
| **Arg-3** | 2.1 | 3.06% | 1.66% | 7.78 |
| **Arg+PFMeUPA-1** | 1.6 | 1.75% | 10.74% | 7.50 |
| **Arg+PFMeUPA-2** | 1.7 | 1.81% | 10.01% | 7.40 |
| **Arg+PFMeUPA-3** | 2.0 | 1.92% | 11.15% | 7.26 |

Note: Initial $NaHCO_3$ added to medium was 20 mM; No initial $H_2$ in the headspace.

### Table S5 Transcriptional regulation of ZM-1 genes in arginine catabolism

| Reaction | Enzyme | Gene ID in ZM-1 | $Log_2$ fold-change in Arg+PFMeUPA vs. Arg | Adj. *p* |
|---|---|---|---|---|
| **1** | Arginine deiminase | 2617 | 1.2[a] | < 0.01[a] |
| **2** | Ornithine carbamoyltransferase | 2616 | 1.2 | < 0.01 |
| **3** | Carbamate kinase | 2615 | 0.8 | < 0.01 |
| **4** | Arginase | 2819 | 1.7 | < 0.01 |
| **5** | Ornithine/alanine racemase | 570 | -0.2 | 0.40 |
| **6** | D-ornithine 4,5-aminomutase E subunit<br>D-ornithine 4,5-aminomutase S subunit | 572<br>573 | -0.3<br>0.06 | 0.19<br>0.80 |
| **7** | 2,4-diaminopentanoate dehydrogenase | 1704<br>1703 | -0.6[b]<br>0.1 | < 0.01<br>0.70 |
| **8** | 2-amino-4-ketopentanoate thiolase, α subunit<br>2-amino-4-ketopentanoate thiolase, β subunit | 1705[c]<br>1706 | 0.5<br>0.1 | 0.15<br>0.74 |
| **9** | Acetate kinase | 3226 | 0.3 | 0.30 |
| **10** | Alanine racemase | 263<br>668 | -0.3<br>0.2 | 0.21<br>0.42 |
| **11-1** | Alanine dehydrogenase | 3381[c] | 1.0 | < 0.01 |
| **11-2** | Alanine aminotransferase | 2446 | 0.1 | 0.5 |
| **12** | Pyruvate:ferredoxin (flavodoxin) oxidoreductase | 2583 | 0.04 | 0.87 |
| **13** | Glutamate dehydrogenase | 2261 | 0.3 | 0.31 |
| **14** | Ornithine cyclodeaminase | 769[c] | 0.7 | < 0.01 |
| **15** | Tungsten-containing aldehyde:ferredoxin oxidoreductase | 1242 | 1.7 | <0.01 |

[a]red: > 1.5-fold increase or adjusted $p \leq 0.01$;
[b]blue: > 1.5-fold decrease;
[c]low transcriptional levels (100 – 500 normalized counts) under both conditions.

### Table S6 Transcriptional regulation of ZM-1 genes related to oxidative energy conservation

| Enzyme | Gene ID in ZM-1 | $Log_2$ fold-change in Arg+PFMeUPA vs. Arg | Adj. *p* |
|---|---|---|---|
| Rnf Complex RnfB | 3726 | 1.2[a] | < 0.01[a] |
| Rnf Complex RnfA | 3727 | 0.8 | < 0.01 |
| Rnf Complex RnfE | 3728 | 0.6 | < 0.01 |
| Rnf Complex RnfG | 3729 | 0.8 | < 0.01 |
| Rnf Complex RnfD | 3730 | 0.3 | 0.14 |
| Rnf Complex RnfC | 3731 | 0.6 | < 0.01 |
| V-type ATP synthase subunit I | 743 | 0.5 | 0.03 |
| V-type ATP synthase subunit K | 744 | 0.4 | 0.17 |
| V-type ATP synthase subunit E | 745 | 0.8 | < 0.01 |
| V-type ATP synthase subunit C | 746 | 0.8 | < 0.01 |
| V-type ATP synthase subunit F | 747 | 1.4 | < 0.01 |
| V-type ATP synthase subunit A | 748 | 0.8 | < 0.01 |
| V-type ATP synthase subunit B | 749 | 0.6 | 0.02 |
| V-type ATP synthase subunit D | 750 | 0.6 | 0.03 |
| Electron-bifurcating [Fe-Fe] hydrogenase (HydA) | 2308 | -2.8 | < 0.01 |
| | 3809 | 0.3 | 0.1 |
| NuoF (HydB) | 2307 | -2.8[b] | < 0.01 |
| | 3810 | 0.6 | 0.01 |
| NuoE (HydC) | 2306 | -2.8 | < 0.01 |
| | 3812 | 1.2 | < 0.01 |
| NuoG (HydD) | 3811 | 0.8 | < 0.01 |
| [Fe-Fe] hydrogenase (HydA) | 2552 | -2.0 | < 0.01 |
| | 3735 | 0.5 | 0.03 |
| Flavocytochrome c (fumarate/urocanate reductase) | 1154 | -1.8 | < 0.01 |
| | 1239 | 0.8 | < 0.01 |
| | 2844[c] | -1.8 | < 0.01 |
| Electron bifurcating acyl-CoA dehydrogenase (NAD+, ferredoxin) | 1289 | -0.4 | 0.01 |
| Electron bifurcating acyl-CoA dehydrogenase, electron transfer flavoprotein beta | 1290 | -0.3 | 0.18 |
| Electron bifurcating acyl-CoA dehydrogenase, electron transfer flavoprotein alpha | 1291 | -0.6 | < 0.01 |
| ferredoxin | 247 | 2.7 | < 0.01 |
| | 487 | 0.3 | 0.39 |
| | 2254 | 0.6 | < 0.01 |
| Fe-S cluster assembly complex SufC | 2404 | 2.2 | < 0.01 |
| Fe-S cluster assembly complex SufB | 2405 | 2.5 | < 0.01 |
| Fe-S cluster assembly complex SufD | 2406 | 2.7 | < 0.01 |
| Fe-S cluster assembly complex SufS | 2407 | 2.8 | < 0.01 |
| Fe-S cluster assembly complex SufU | 2408 | 2.6 | < 0.01 |
| Fe-S oxidoreductase (Radical SAM protein) | 2892 | 6.3 | <0.01 |

[a]red: > 1.5-fold increase or adjusted $p \leq 0.01$;
[b]blue: > 1.5-fold decrease;
[c]low transcriptional levels (100 – 500 normalized counts) under both conditions.

**Table S7** Transcriptional regulation of ZM-1 genes involved in gluconeogenesis

| Enzyme | Gene ID in ZM-1 | $Log_2$ fold-change in Arg+PFMeUPA vs. Arg | Adj. *p* |
|---|---|---|---|
| Phosphoenolpyruvate carboxykinase [GTP] | 1772 | 6.0[a] | < 0.01[a] |
| Pyruvate, phosphate dikinase | 3958 | 1.0 | < 0.01 |
| Enolase | 346 | 0.8 | < 0.01 |
| 2,3-bisphosphoglycerate-independent phosphoglycerate mutase | 347 | 0.6 | < 0.01 |
| Phosphoglycerate kinase | 349 | 1.2 | < 0.01 |
| NAD-dependent glyceraldehyde-3-phosphate dehydrogenase (GAPDH) | 350 | 1.8 | < 0.01 |
| Fructose-bisphosphate aldolase class II | 1076 | 2.9 | < 0.01 |
| Fructose-1,6-bisphosphatase | 1372 | 1.3 | < 0.01 |

[a]red: > 1.5-fold increase or adjusted $p \leq 0.01$.

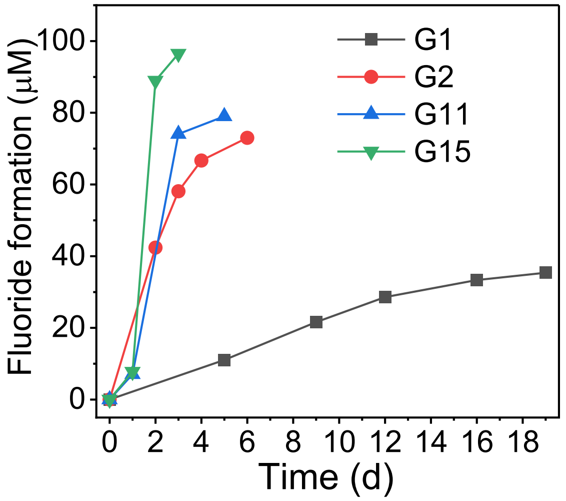


**Figure S1**. Defluorination activity of enrichment cultures over the enrichment process (G1, 2, 11, and 15 are enrichment cultures after 1, 2, 11, and 15 times of subculturing using the same enrichment conditions: 35 mM lactate and 100 µM PFMeUPA).

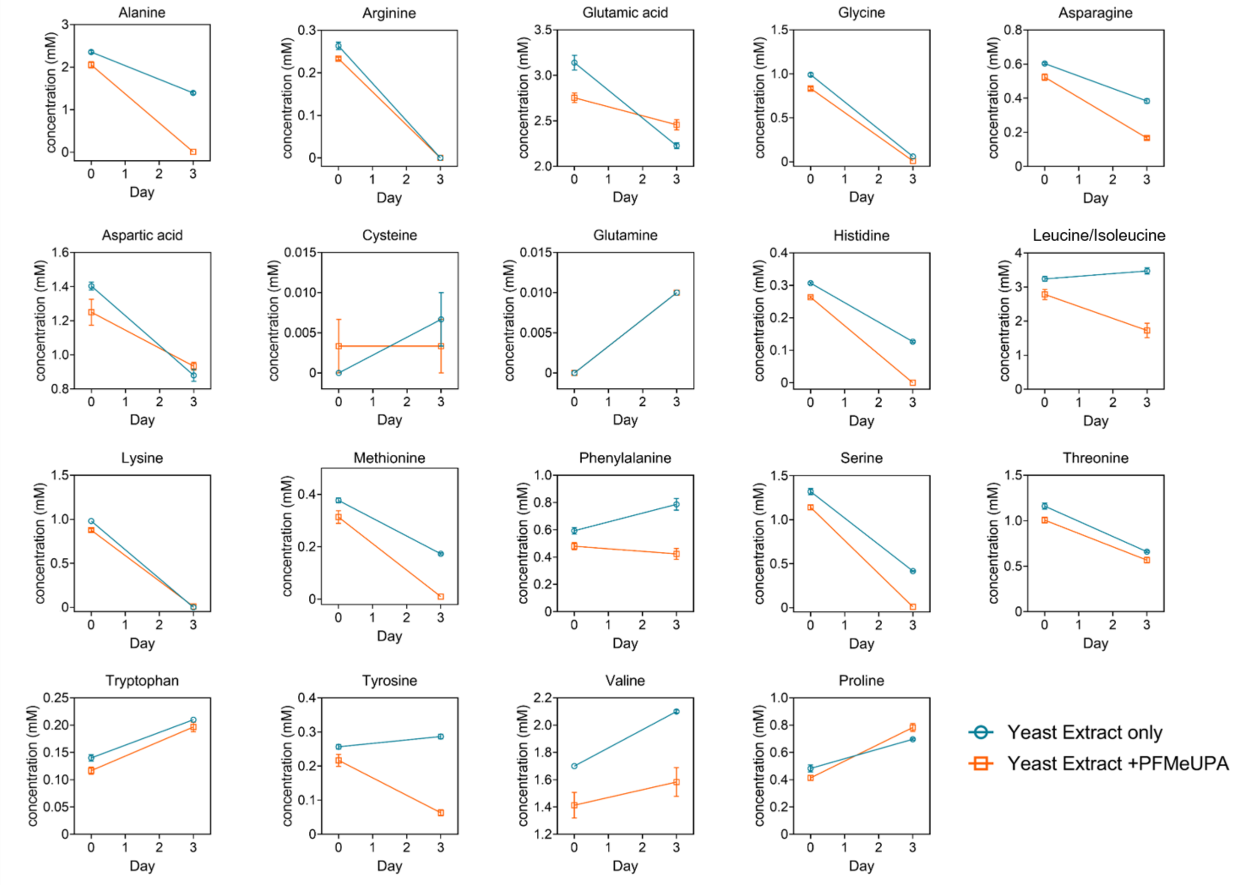


**Figure S2**. Amino acid utilization in ZM-1 grown on yeast extract (0.3% w/v) only and with PFMeUPA (5 mM).

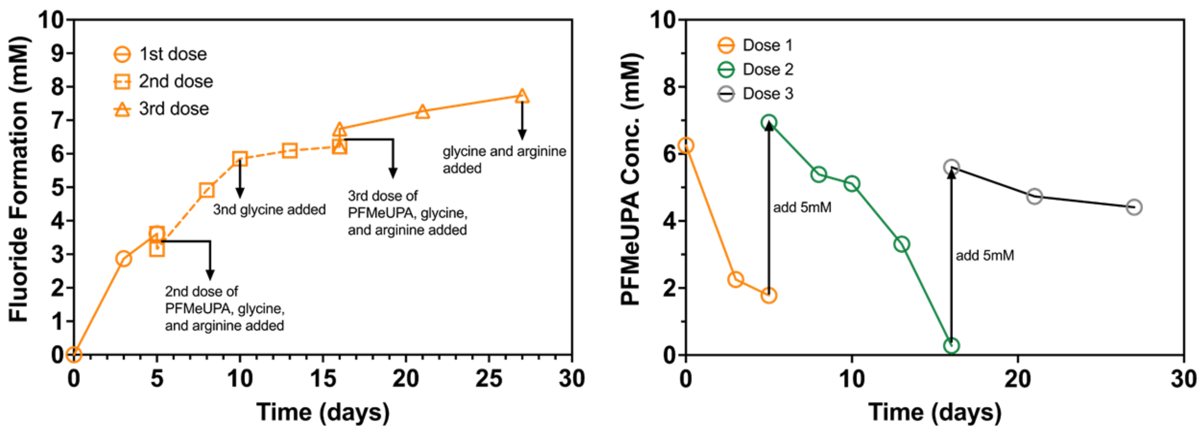


**Figure S3**. Multiple-dose PFMeUPA defluorination by ZM-1 (each dose of PFMeUPA: 5 mM; arginine and glycine addition: 10 mM each).

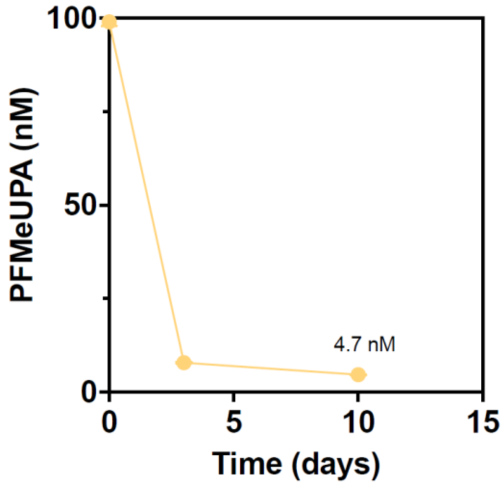


**Figure S4**. Defluorination of 100 nM PFMeUPA by ZM-1.

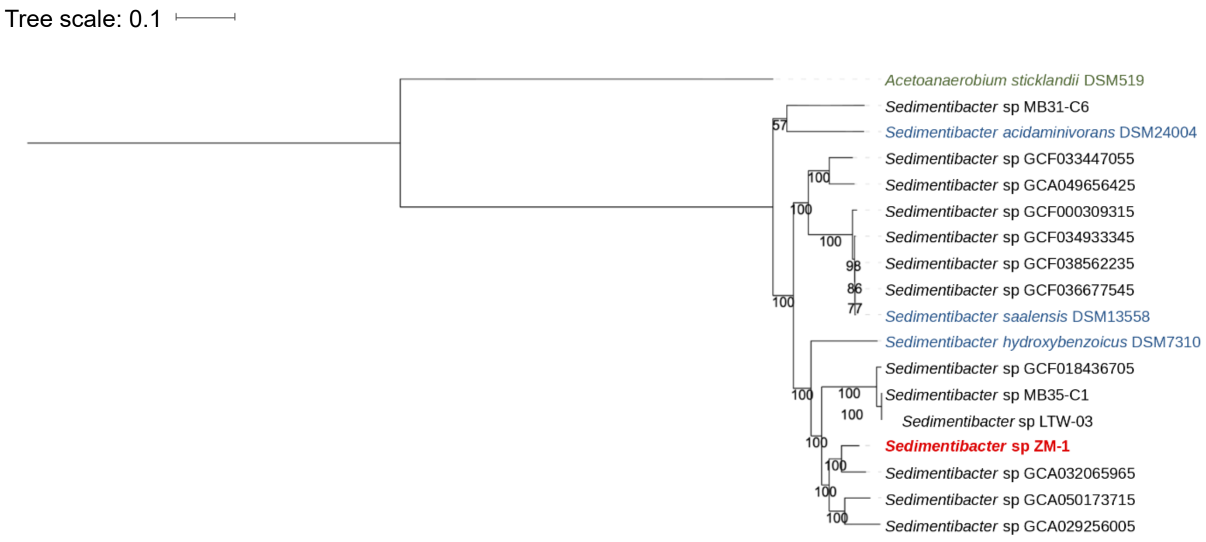


**Figure S5**. Phylogenetic tree of *Sedimentibacter sp.* ZM-1 (Outside species: *Acetoanaerobium sticklandii* DSM519; three *Sedimentibacter* isolates purchased from DSMZ and investigated in this study are highlighted in blue; ZM-1 isolated in this study is highlighted in red).

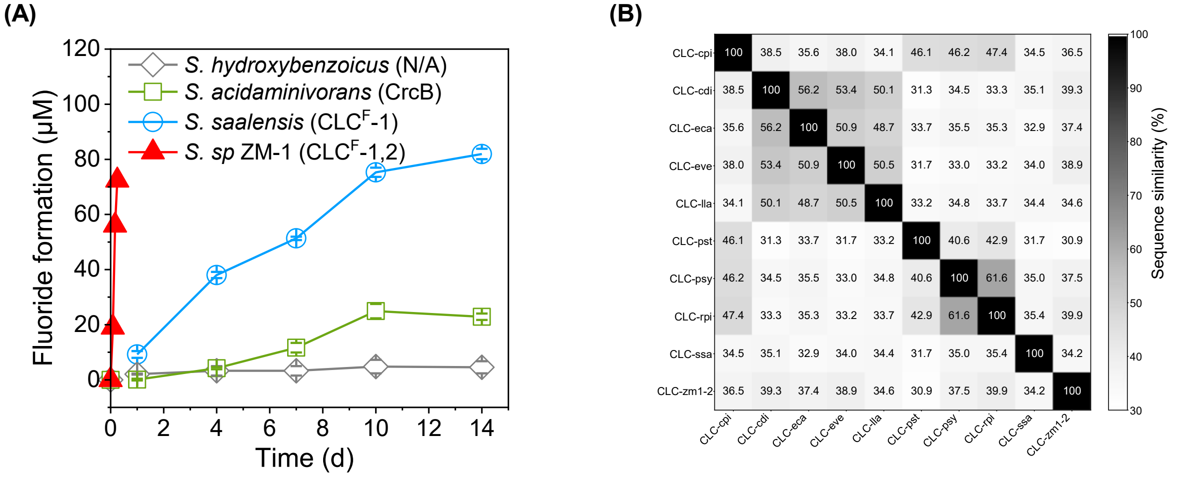


**Figure S6**. PFMeUPA defluorination and fluoride ion exporter comparison (A: PFMeUPA defluorination by different *Sedimentibacter* species, in parentheses are fluoride exporters identified in the genome, n = 3; B: pairwise similarity of $CLC^F$ homologues, including eight previously studied ones (Ref 21 in the main text) with NCBI accession numbers from top to bottom YP_003123680.1, YP_001087131.1, ZP_05647488.1, ZP_02027167.1, YP_001032764.1, YP_003370005.1, ZP_07250577.1, and YP_001892513.1; CLC-ssa: the $CLC^F$ homologue in *S. saalensis*, NCBI accession number WP_145080895.1; CLC-zm1-2: $CLC^F$-2 in ZM-1 genome, GeneID in the draft genome: 1445, Table S3).

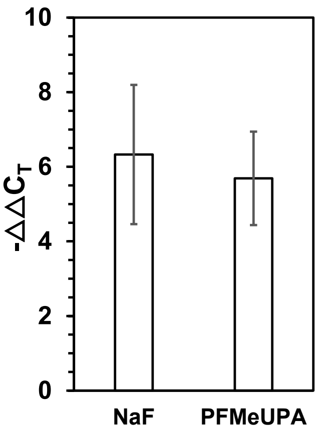


**Figure S7**. Differential gene expression of the *eriC* gene encoding CLC[F]-2 in ZM-1 culture grown on arginine and glycine exposed to 3 mM NaF and in ZM-1 culture grown on arginine and glycine with 3 mM fluoride formation from 5 mM PFMeUPA addition after two days of incubation (*gyrB* was used as the reference housekeeping gene for the $\Delta\Delta C_T$ calculation; n = 3).

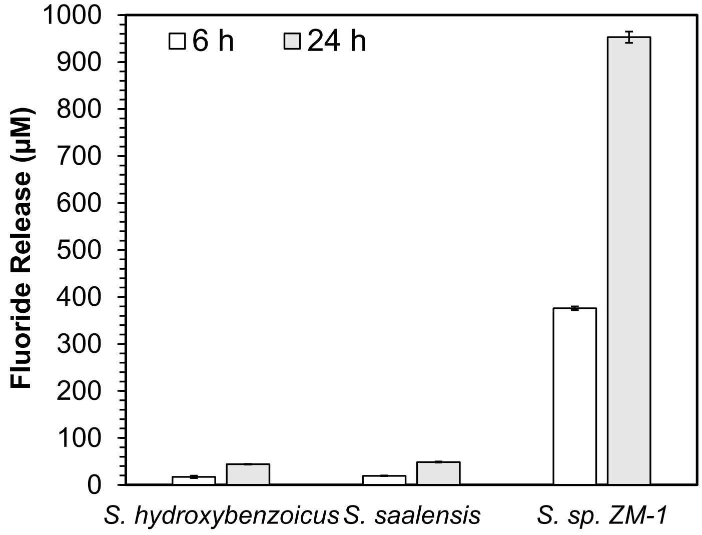


**Figure S8**. Fluoride formation from 1 mM PFMeUPA in *in vitro* assays (1 mL) using crude extracts from *S. hydroxybenzoicus*, *S. saalensis*, and *S. sp* ZM-1 under anaerobic conditions (n = 3; Note: crude extracts were obtained from 100 mL pre-grown cells, OD600 = 1.2, which were lysed by sonication in 20 mL lysis buffer solution containing 50 mM Tris-HCl (pH 8.0), 50 mM NaCl, 5% (v/v) glycerol, and 1 mM 1,4-dithiothreitol. The supernatant after lysis and centrifugation (14,000 × g, 4°C for 45 min) was used for the *in vitro* assay).

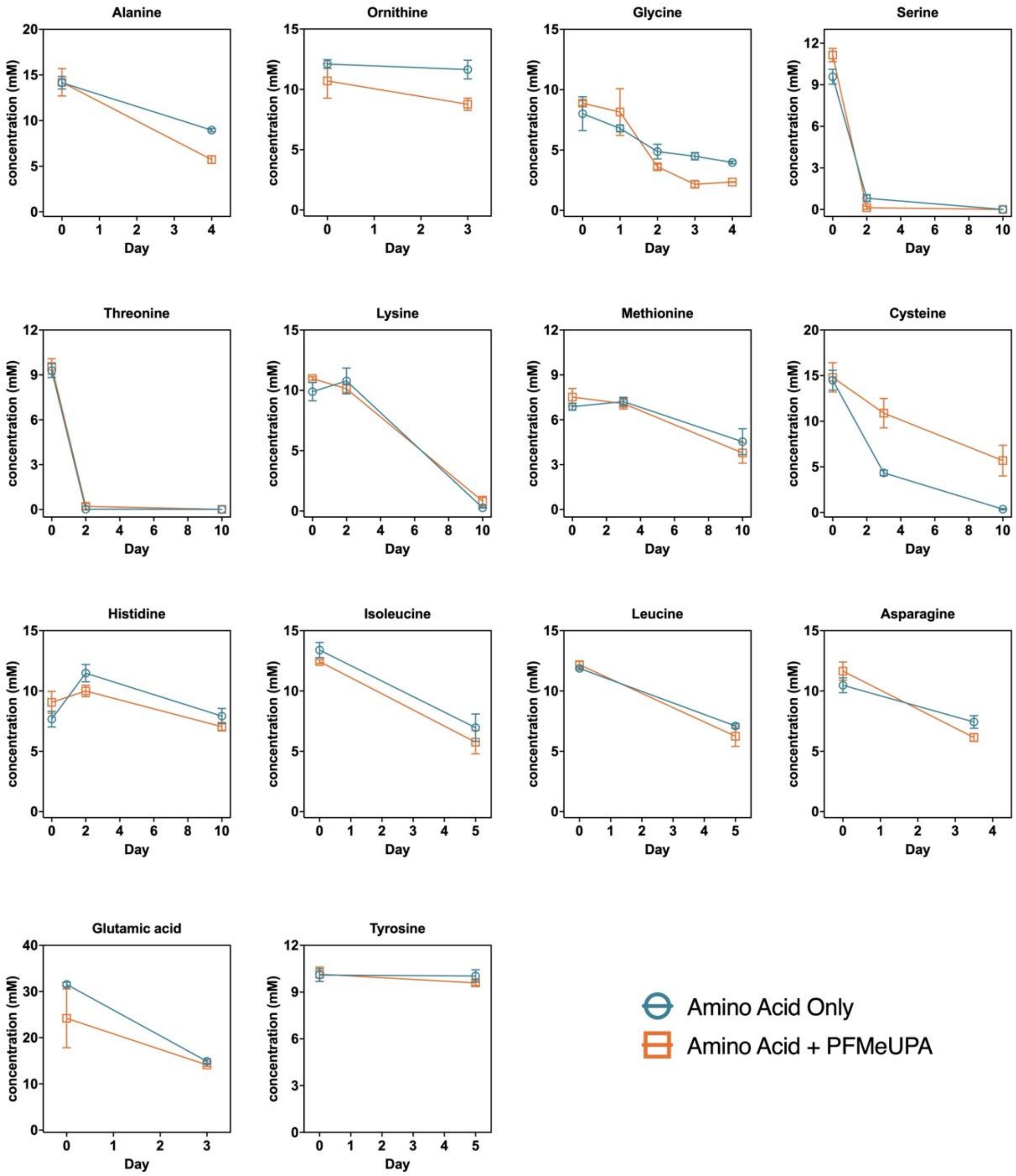


**Figure S9**. Utilization of individual amino acids (10 mM) by ZM-1 grown in basal medium containing 0.3% (w/v) yeast extract with and without PFMeUPA (5 mM) (amino acids contained in the yeast extract and shown to be consumed in **Fig. S2** were included).

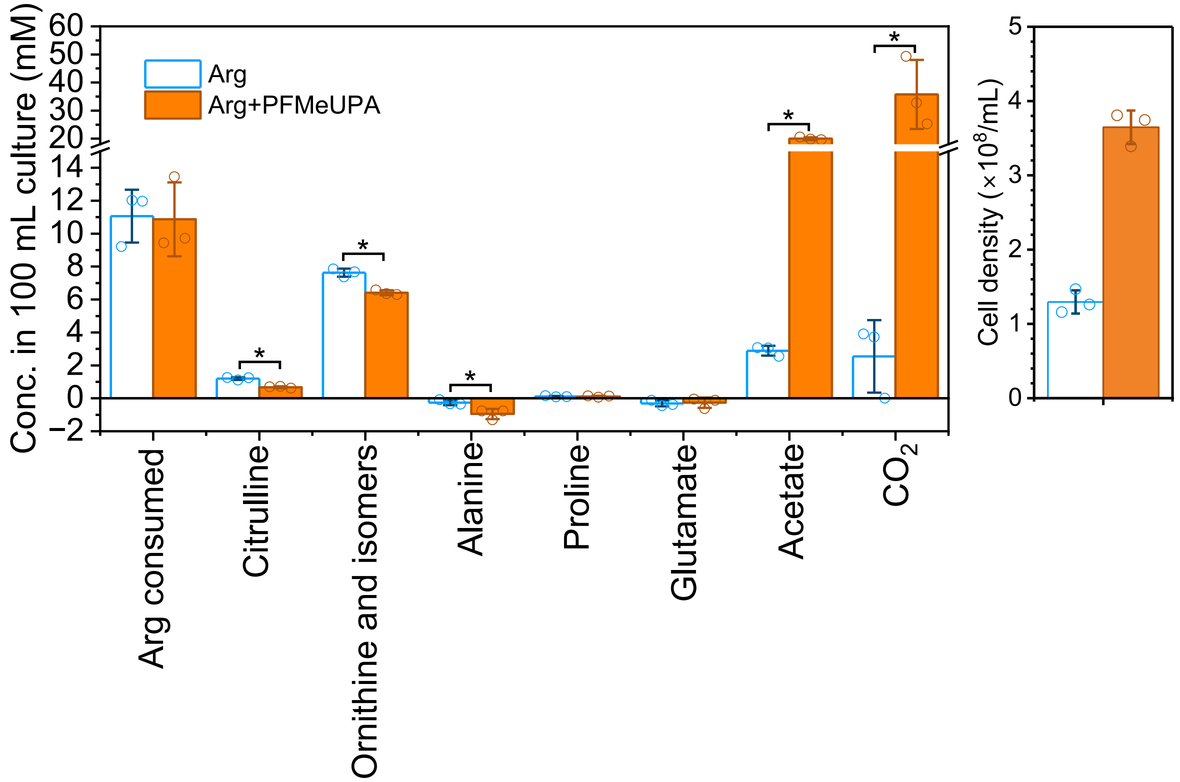


**Figure S10**. Arginine consumption, metabolite formation, and cell growth of ZM-1 grown on arginine (10 mM) with and without PFMeUPA (5 mM) at the end of incubation (*: Student's t-test, $p < 0.05$, n = 3; results are in the same batch as those shown in **Fig. 3C** in the main text).
